# RESIC: A tool for comprehensive adenosine to inosine RNA Editing Site Identification and Classification

**DOI:** 10.1101/2021.04.11.439401

**Authors:** Dean Light, Roni Haas, Mahmoud Yazbak, Tal Elfand, Tal Blau, Ayelet T. Lamm

**Affiliations:** Faculty of Biology, Technion- Israel Institute of Technology, Technion City, Haifa 32000, Israel

**Keywords:** SARS-CoV-2, ADAR, epitranscriptome, hyper-editing, Interferon

## Abstract

Adenosine to inosine (A-to-I) RNA editing, the most prevalent type of RNA editing in metazoans, is carried out by adenosine deaminases (ADARs) in double-stranded RNA regions. Several computational approaches have been recently developed to identify A-to-I RNA editing sites from sequencing data, each addressing a particular issue. Here we present RESIC, an efficient pipeline that combines several approaches for the detection and classification of RNA editing sites. The pipeline can be used for all organisms and can use any number of RNA-sequencing datasets as input. RESIC provides 1. The detection of editing sites in both repetitive and non-repetitive genomic regions; 2. The identification of hyper-edited regions; 3. Optional exclusion of polymorphism sites to increase reliability, based on DNA, and ADAR-mutant RNA sequencing datasets, or SNP databases. We demonstrate the utility of RESIC by applying it to human, successfully overlapping and extending the list of known putative editing sites. We further tested changes in the patterns of A-to-I RNA editing, and RNA abundance of ADAR enzymes, following SARS-CoV-2 infection in human cell lines. Our results suggest that upon SARS-CoV-2 infection, compared to mock, the number of hyper editing sites is increased, and in agreement, the activity of ADAR1, which catalyzes hyper-editing, is enhanced. These results imply the involvement of A-to-I RNA editing in conceiving the unpredicted phenotype of COVID-19 disease. RESIC code is open-source and is easily extendable.

## Introduction

The conversion of adenosine to inosine (A-to-I) in double-stranded RNA regions, by adenosine deaminases (ADARs) enzymes, is the most common form of RNA editing in metazoans (Bazak et al., 2014). This type of RNA editing is crucial for normal development of an organism and has a major role in the innate immune response (Mannion et al., 2014; Ganem and Lamm, 2017; Eisenberg and Levanon, 2018). It was shown that changes in editing events are correlated with several types of diseases; among them is cancer (Maas et al., 2006; Galeano et al., 2012; Gallo and Locatelli, 2012; Kung et al., 2018). Editing sites may serve as biomarkers for cancer and ADAR enzymes are considered as promising gene therapy agents to fight cancer (Ganem et al., 2017). In addition, ADARs are known to be involved in regulation of innate immune response by blocking the interferon response upon viral infection (Quin et al., 2021). For these reasons, A-to-I RNA editing is an extensively studied research field in many organisms, and identification of editing sites is of major interest.

In recent years, many efforts have been invested in developing computational approaches to detect A-to-I RNA editing sites from sequencing data (Pinto and Levanon, 2019). Since inosine is very similar in structure to guanosine, inosine is interpreted as guanosine by polymerases during sequencing. This enables the detection of editing sites by comparing between DNA and RNA sequences, to track adenosine to guanosine (A-to-G) mismatches. However, the detection should be carefully performed to avoid false reports due to sequencing and alignment mistakes, alterations in sequence originated from polymorphism, somatic mutations, or other changes which are not the result of editing events (Pinto and Levanon, 2019). The problem is exacerbated by the fact that editing in humans frequently occurs in repetitive regions (Athanasiadis et al., 2004; Blow et al., 2004; Kim et al., 2004; Levanon et al., 2004; Levanon et al., 2005; Barak et al., 2009; Kleinberger and Eisenberg, 2010; Osenberg et al., 2010; Paz-Yaacov et al., 2010; Wu et al., 2011b), which tend to cause alignment errors (Treangen and Salzberg, 2011).

Several tools developed to detect A-to-I RNA editing sites are based on comparison between RNA-seq reads and DNA reference genome. Among these tools are REDItools, which suggest simple comparison using samtools (Picardi and Pesole, 2013), and GIREMI that focused on distinguishing between SNPs and editing, relying on existing SNP databases and a given RNA-seq data (Zhang and Xiao, 2015). Some tools support a direct comparison between RNA-seq reads and DNA reads from the same source, allowing editing site identification without the need for previous knowledge (Picardi and Pesole, 2013; Lee et al., 2015; Wang et al., 2016; Piechotta et al., 2017). A major advantage in comparing between DNA and RNA sequences of the same biological sample is the ability to increase accuracy by excluding changes deriving from unpublished SNPs (Pinto and Levanon, 2019). Another way to increase the results accuracy is parallel comparison between several RNA-seq datasets of several individuals, while taking into consideration that true editing sites would appear in all or most samples (Ramaswami et al., 2013; Wang et al., 2016; Goldstein et al., 2017).

Hyper-editing by ADAR enzymes, which is defined as multiple A-to-I editing sites in a proximity, is a widespread phenomenon. Since most tools designed to identify editing sites are based on the detection of a limited number of mismatches in read alignments (to reduce alignment errors and running time), hyper-editing events, which result in multiple mismatches in a single read, are usually unexposed. Therefore, several recent methods were specially oriented to track hyper-editing sites. Wu et al. 2011 (Wu et al., 2011a) and Porath et al. 2014 (Porath et al., 2014) developed methods that are based on the conversion of unmapped read-sequences to a three-base code genome and thus enable identification of hyper-editing sites. Namely, all As are transformed to Gs in the reference genome and in the RNA-seq reads that previously failed to align, and realignment is then carried out. Following reversion to original sequences, hyper-editing sites, which are rich with A-to-G mismatches, can be located. In both studies, conversion to a three-base code was repeated for all possible nucleotide pairs. It was shown that A-to-G editing was enriched over the other editing types.

Despite the efforts to develop computational tools for A-to-I RNA editing site detection from sequencing data, to date there is not a single platform enabling robust detection of editing sites of different classes and their classification. Here we present RESIC, which enables detection and classification of A-to-I RNA editing sites of different types in a single tool. We expanded the pipeline we previously applied to identify editing sites in repetitive and non-repetitive regions (Goldstein et al., 2017) and adopted the method by Wu et al. 2011 (Wu et al., 2011a) and Porath et al. 2014 (Porath et al., 2014) to find hyper-editing sites. The tool includes an alignment-graph of distinctive architecture and several filtration steps to reduce false identifications. RESIC also enables distinguishing between polymorphism and editing events to increase readability, by using DNA sequences, ADAR mutant RNA-sequencing datasets, or a SNP database. We demonstrate the utility of RESIC by applying it to mapping A-to-I RNA editing sites in 16 human tissues, from the Illumina Human Body Map project, analyzed for a similar purpose by others (Zhu et al., 2013; Bazak et al., 2014; Porath et al., 2014). Our analysis reproduced known putative editing sites, detected by others and included in the RADAR database (Ramaswami and Li, 2014), and extended the list of known sites.

Since aberrant Interferon (IFN) and cytokine responses were observed in COVID-19 patients (Moore and June, 2020) and ADAR1 was shown to activate the interferon reaction (Baños-Lara et al., 2013), we further interrogate the activity of A-to-I RNA editing upon SARS-CoV-2 infection. We show that in SARS-CoV-2 infected samples, compared to mock, ADAR1 is the only A-to-I RNA editing enzyme that is differentially expressed, and the numbers of A-to-I hyper editing sites are larger.

## Methods and definitions

### RESIC Algorithmic definitions

RESIC enables the user to supply DNA or RNA datasets that should exhibit the desired editing phenomena and DNA or RNA sequencing datasets that should not exhibit the desired editing phenomena. The latter group is used to exclude changes deriving from SNPs. Since nucleotide changes in the former sequencing datasets correspond to positive evidence of that sites undergoing editing and the latter datasets correspond to negative evidence, we term these sets of datasets as positive and negative datasets. RESIC is completely reference agnostic; the user provides whichever reference file they wish to use for the alignment as well.

#### Ambiguous read filtering

For ambiguous read filtering, we adopted the method of (Porath et al., 2014). Briefly, we filtered out the reads that meet the next criteria: one or more nucleotides represent over 60% or under 10% of the read sequence, more than 10% Ns (when a base call could not be done), average Phred quality score < 25, and more than 20 repeats of a single nucleotide in a row.

#### Alignment scheme

We define an alignment scheme to be a 4-tuple *S* = (*A, p, f*_1_, *f*_2_) where *A* is an alignment algorithm, *p* is a list of alignment parameters for *A, f*_1_ is a preprocessing function of the raw datasets and *f*_2_ is a postprocessing function for aligned and misaligned reads. These seemingly verbose definitions enable RESIC to decouple the choice of alignment algorithm from the rest of the modules in RESIC.

Let *S* be an alignment scheme, *L* be a sequencing dataset and *R* be a genome reference, we define *P*_*S,L,R*_ and *N*_*S,L,R*_to be the aligned and misaligned read fractions resulting from running S on L and R. We define S(*L, R*) = (*P*_*S,L,R*_, *N*_*S,L,R*_).

We say that a scheme is normal if *f*_1_ and *f*_2_ are identity functions in said scheme. Pseudo code for calculating S(L,R):

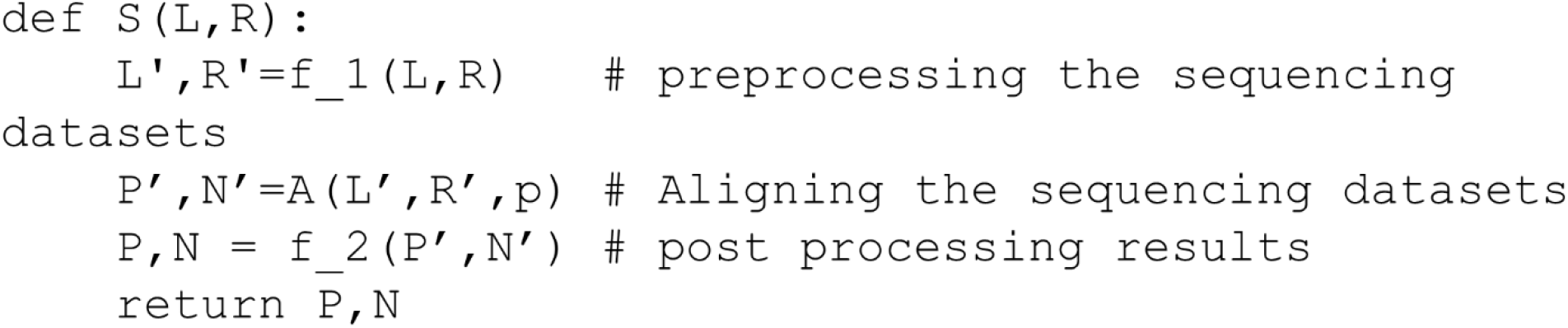

#### Graph Aligner

Given a directed acyclic graph *G* = (*V, E*) where nodes in *V* are alignment schemes, *L* a sequencing dataset and *R* a reference we define new alignment schemes *G*_*v*_(*L, R*) for each *v* ∈ *V* to be defined as follows:

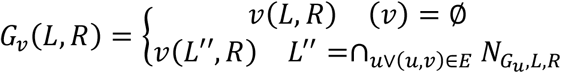

We define *G*(*L, R*) to be a set of aligned sequencing datasets 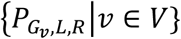. Simply put, we perform the alignment scheme of node v on all read fragments that were misaligned in any of v’s ancestors.

#### 3nt genome alignment scheme

Let *X* and *Y* be two distinct nucleotides. To be able mapping hyper editing sites, we apply the 3nt alignment scheme by which each appearance of either *X* or *Y* is transformed into | in both the sequencing datasets (reads) and the reference genomes. That was similarly done by others (Wu et al., 2011a; Porath et al., 2014). However, we present an advanced 3nt technique to map hyper antisense reads as was not described elsewhere, to the best of our knowledge. First, for each *X* and *Y* nucleotides pairs, we first apply the scheme to the reads at the given node (see graph aligner) and to the reference sense strand. Next, in order to identify hyper editing sites on the antisense strand, for each *X* and *Y* nucleotides pairs, we create the complement reference genome, based on the original reference, and reverse the reads that were unmapped in the previous step, to achieve the 3’ -5’ direction, same as the created reference. Then we reapply the 3nt alignment scheme while considering the manipulation when recording the mapped reads as aligned to the antisense.

In each step, after mapping the reads, aligned and unaligned reads are reverted to their original sequence via custom python scripts. **Figure S1** illustrates into details the 3nt genome alignment scheme. To conduct the 3nt scheme we use awk (Aho et al., 1996) and sed (Free Software Foundation).

#### Site filtering

After performing the graph alignment for each of the given sequencing datasets, samtools (Li et al., 2009) is used to convert the files into pileup format. Then, several filtering steps are performed as detailed below. All parameters (*l, k1, k2, u, r*, and *c*) are user defined. First, sites with no nucleotide changes and sites covered by less than *l* reads are discarded. We discarded sites from the positive datasets if those same sites appeared in any negative dataset with a nucleotide change.

#### Editing percent filtering

For each positive sequencing dataset, we filter out any site: (1) whose most abundant nucleotide change constitutes less than *k*_1_ percent or more than *k*_2_ percent of the reads mapped to that site (2) whose other nucleotide changes constitute over *u* percent of the reads mapped to that site (3) whose most abundant nucleotide change is in at least *r* reads. We term: *k*_1_, the minimal editing percent threshold, *k*_2_, the maximal editing percent threshold, *r*, the editing read threshold, and *u*, the editing noise threshold.

#### Unique site filtering

We filter all sites that were defined as editing sites at the previous step, under more than one editing category (e.g. non-repetitive and Hyper A to C), if they represented more than one type of nucleotide change (e.g. once A to G and the other time A to C).

#### Hyper editing filtering

Deriving from our method, it may be possible that under the hyper editing categories, a non-hyper editing site would be recorded. Namely, for each pair of nucleotides X and Y that we perform the 3nt genome scheme, other nucleotide mismatches than hyper X to Y or Y to X may be noted, enabled by the new conditions created by the 3nt scheme. Therefore, we filter the hyper editing files to include only X to Y or Y to X changes (see an illustration in **Figure S1**).

#### Consensus

We filter out any sites that are not present in over *c* percent of positive datasets.

### A-to-I editing identification pipe

We implemented a hyper editing alignment scheme and built an alignment graph that could target any editing type. Specifically, in the analysis described here we only applied RESIC to A-to-I RNA editing.

#### A-to-I RNA editing alignment graph

In our screen for A-to-I editing sites, we define two classes of alignment schemes, non-repetitive alignment for reads that map uniquely to the genome and repetitive for repetitive regions or regions that cannot be differentiated by our reads via alignment. Our graph alignment, summarized in **Figure S2**, is as follows: We align sequencing datasets using the non-repetitive normal scheme followed by the repetitive normal scheme. Then we branch out and for each pair of distinct nucleotides X and Y, we perform the non-repetitive 3nt genome scheme, and the repetitive 3nt genome scheme.

#### RNA editing profiling of Illumina bodyMap2 transcriptome

RNA-seq datasets from 16 human tissues (Illumina Human Body Map 2.0 Project; GEO accession number GSE30611) that were sequenced at 75 single read (SR), were downloaded from SRA. FastQC was used to control the read quality and trimming was performed accordingly. Reads were further collapsed and then taken for RESIC run. For the underline sequencing algorithm, we used Bowtie (Langmead et al., 2009) alignment tool. For the non-repetitive and repetitive alignments, we configured bowtie to align to fragments if they map to under 2, or 20 different genomic locations, respectively, with at most 3 single base mismatches and to consider matches for a read r as the set of alignment results for r with the smallest alignment score (-m 2 -n 3 --best --strata, -l 50 --chunkmbs 200, and -m 20 -n 3 --best --strata -l 50 --chunkmbs 200, respectively). Similar alignment was used in Goldstein et al. (Goldstein et al., 2017). For the site filtration steps, we choose *l* = 2 to be the coverage per site threshold, *k*_1_ = 30 and *k*_2_ = 99 for the editing minimal and maximal percent threshold, respectively, u = 3 for the noise thresholds, r =2 for the editing read threshold, and *c* = 0; for consensus threshold. The site lists obtained for each tissue were filtered to have only A-to-I sites, namely A-to-G, or T-to-C mismatches in both strands.

The list of obtained editing sites was compared to the entire list from RADAR database (Ramaswami and Li, 2014), We considered as shared editing sites, sites that are included in the RADAR list or sites that have gene annotations similar to the ones appeared in the RADAR list.

#### Profiling of SARS-CoV-2 infected Calu-3 cells

Raw RNA-seq data of Calu-3 human Lungadenocarcinoma cells infected with SARS-CoV-2 virus or mock, were downloaded from SRA, BioProject PRJNA615032. FastQC was used to control the read quality and trimming was performed accordingly. Reads were collapsed and first aligned to the SARS-CoV-2 reference genome version NC_045512.2 using bowtie. Alignment to the SARS-CoV-2 genome was made to exclude reads that are originated from the virus for further analysis, and to validate that in contrast to the mock samples, the SARS-CoV-2 samples are infected with the virus. Indeed, few thousands of reads were mapped to the SARS-CoV-2 genome, only for the SARS-CoV-2 infected samples. We applied RESIC separately on the unaligned reads of the mock and SARS-CoV-2 infected samples (3 biological replicates each) to identify changes in RNA editing events upon coronavirus infection. For the underline sequencing algorithm, we used Bowtie (Langmead et al., 2009) alignment tool. For the non-repetitive and repetitive alignments, we configured bowtie to align to fragments if they map to under 2, or 100 different genomic locations, respectively, with at most 3 single base mismatches (-m 2 -n 3 --best --strata, -l 50 --chunkmbs 200, and -m 100 -n 3 --best --strata -l 50 --chunkmbs 200, respectively). For the site filtration steps, we choose *l* = 2 to be the coverage per site threshold, *k*_1_ = 30 and *k*_2_ = 99 for the editing minimal and maximal percent threshold, respectively, u = 3 for the noise thresholds, and r =2 for the editing read threshold. The consensus module was run with *c* = 0.5. We then filtered the site lists obtained to have only A-to-I sites. Since the RNA library preparation strategy was stranded (the sequenced strand must be from the actual expressed strand), we filtered the files obtained to include only A-to-G, and not T-to-C sites.

To perform differential expression analysis, we mapped the same unaligned reads that were used for RESIC analysis before, to the human transcriptome version GRCh37 (hg19) using bowtie. Gene expression levels were evaluated by read counts. We then compared our created gene counts to the already processed counts downloaded from GEO: GSE147507. Although read count values were not identical, as expected due to the use of different alignment tools, the trend was the same.

We performed differential expression analysis (DEA), using DESeq2 (Love et al., 2014), with lfcShrink function and apeglm shrinkage estimator type.

## Results and discussion

### RESIC – A comprehensive tool for identification RNA editing sites

To have a complete identification of RNA editing sites, which include sites in non-repetitive regions, sites in repetitive regions, and sites in hyper-editing regions, we generated a novel tool termed “RNA Editing Sites Identification and Classification” (RESIC). RESIC composed of an enhanced alignment graph model to identify and classify editing sites by their type, multiple-step filtering process to increase result reliability in a flexible manner (**Figure 1 A-B**) and plots for data visualization (see an example in **Figure 2**).

**Figure 1:**
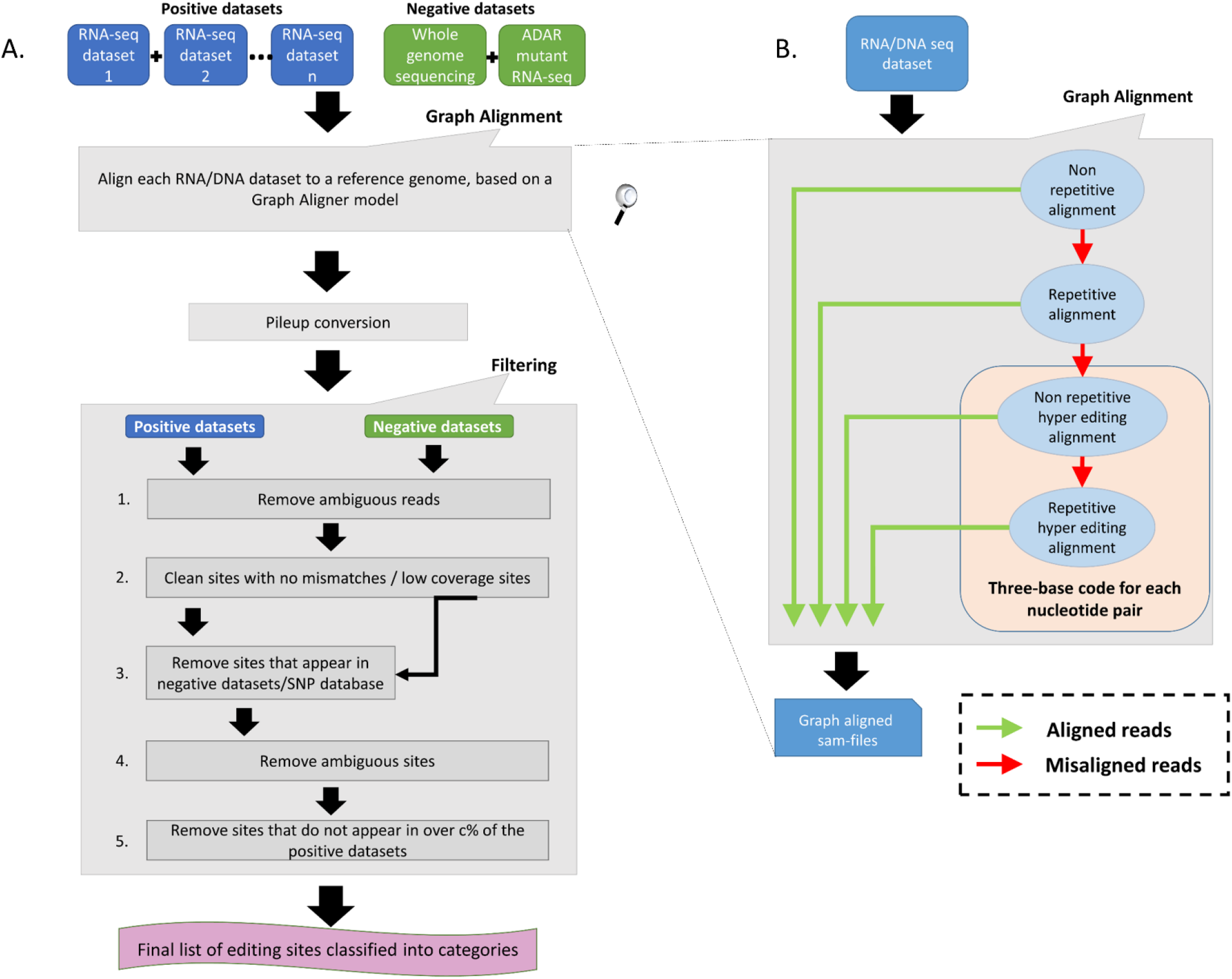
RESIC’s schematic view. (A) Overall description of the RESIC pipeline. First all given sequencing datasets are filtered for ambiguous reads and go through the graph alignment scheme to detect A-to-I editing sites of different classes. The RNA-seq datasets intended for editing-sites identification are termed positive datasets, and the RNA and/or DNA sequencing datasets used to contradict editing-site existence are termed negative datasets. Sam-files for each alignment node are created using a sequence aligner (Langmead et al., 2009) and converted into pileup files using samtools (Li et al., 2009). The next stage includes several filtering steps for removing: 1) sites with no changes compared to the reference or low coverage sites 2) SNPs or mismatches that are not originated in RNA editing, based on comparison to the negative datasets or/and a SNP database (optional). 3) sites with more than one prominent mismatch (large noise) or with low change ratio. 4) sites that do not appear in over c% samples (optional). Finally, a list of editing sites divided into classes are given as an output with descriptive plots. (B) Zoom in on the data flow illustration of the graph aligner model for the four-layer graph used in the study.

To use RESIC, the user should supply positive datasets, i.e. RNA-/DNA sequencing datasets that should exhibit the desired editing phenomena, and a reference genome. The user may also supply negative datasets, DNA or RNA sequencing datasets that should not exhibit the desired editing phenomena, or a SNP database, to contradict editing-site existence.

All datasets are first processed according to the graph alignment (**Figure 1 A-B**). The graph alignment was designed to track editing sites of different types by aligning a given set of reads to the reference genome in a specific parameter configuration setup that represents each editing class. Namely, we demanded unique or multiple alignment to detect nonrepetitive and repetitive sites respectively, Next for sequences that did not align, we converted read-sequences to a three-base code to detect hyper editing sites for all possible nucleotide pairs, as described by Wu et al. 2011 (Wu et al., 2011a) and Porath et al. 2014 (Porath et al., 2014). Our alignment-graph distinctive architecture enables the fluent utilization of an unmapped read-set that was discarded in one alignment level for defining editing sites of a different class in the next level (**Figure 1B**). This enables the identification of multiple editing-site classes in a single platform. While RESIC was built to provide a way to consolidate the many ongoing efforts at A-to-I editing site identification, our graph aligner model is general and robust enough to stand on its own and contribute to general identification of nucleotide changes.

Following alignment, the candidate editing sites that were identified are going throw strict multi-stage filtering process (**Figure 1A**). The filtering process aim is to increase the result reliability considering different types of possible errors. In one type, sites in which there is more than one mismatch type or sites showing low change ratio are suspicious as technical errors likely to be formed during sequencing or alignment and discarded due to low reliability. The user may easily modify the limiting thresholds controlling these filtering steps (i.e. minimal coverage per site, minimal change ratio and maximal noise ratio). In another type, incorrect recognition of SNPs as editing sites can be prevented by excluding sites that show the same nucleotide alterations in both the DNA and the RNA sequences. The user may choose (but it is not mandatory) supplying DNA sequencing data of the same individual used to detect editing sites, for enabling the described DNA based exclusion. Another way in which SNPs can be distinguished from editing sites is parallel comparison between several samples of different individuals to test for the consensus level of editing sites. The rationale behind parallel comparison of various individuals is that true editing sites would appear in all or most samples (Ramaswami et al., 2013; Wang et al., 2016; Goldstein et al., 2017). In addition, biological replicas can eliminate changes that occurred because of sequencing errors. Testing for consensus in editing sites among several samples is a less favorable option to eliminate SNPs, in case DNA sequencing dataset is supplied. The user may choose to neutralize the consensus filtering step or modify the consensus threshold.

### RESIC enhanced the number of identified A-to-I editing sites in *human* tissues

In order to test the utility of the tool, we used RNA-seq datasets from 7 human tissues: adipose, adrenal, brain, breast, colon, kidney, and heart (Illumina Human Body Map 2.0 Project; GEO accession number GSE30611) that were sequenced at 75 single read (SR). These datasets were previously screened for editing sites by others (Zhu et al., 2013; Bazak et al., 2014; Porath et al., 2014).

We used the latest GRCh37 SNP database (NCBI) to eliminate changes that are not originated from A-to-I RNA editing, but from genomic polymorphism. All datasets were processed according to the graph alignment and went through all filtration steps (for parameters setup, see Methods). Since each of the 16 samples is originated from a different tissue, and editing sites may be tissue specific (Picardi et al., 2015), we defined *c* = 0 for consensus threshold. To test the power of RESIC to specifically identify A-to-I editing sites, we compared the output of RESIC (**Supplementary Table S1)** to the collection of A-to-I RNA editing sites, taken from RADAR (Ramaswami and Li, 2014). It is indicated by our comparison (**Table 1**) that over 75% of the non-repetitive sites RESIC identified, and over 65% of non-repetitive hyper sites are also included in the RADAR collection. This large overlap is expected, since RADAR is based, among others, on the same samples analyzed by us, and at the same time strengthening the reliability of RESIC. Since hyper editing sites are less frequently found by traditional tools, dictated by the parameter setup (Pinto and Levanon, 2019) and tools that aimed for tracking hyper-editing sites (Wu et al., 2011a; Porath et al., 2014) are less abundant, it is not surprising that a smaller overlap was obtained for non-repetitive hyper sites, compared to non-repetitive.

**Table 1:**
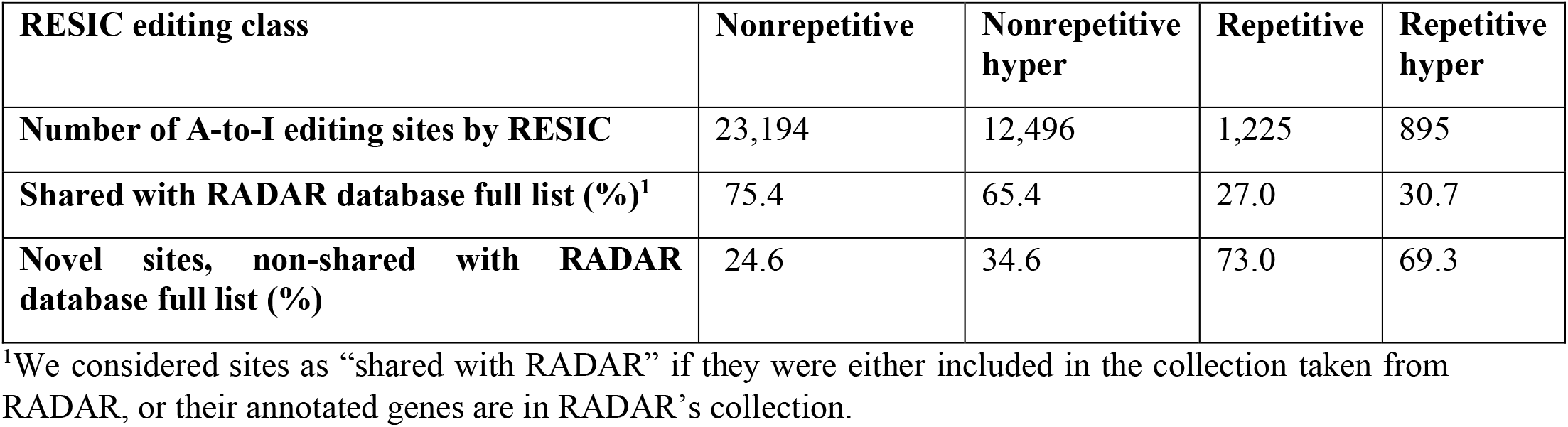
overlap levels of the detected A-to-I editing sites with RADAR database.

| RESIC editing class | Nonrepetitive | Nonrepetitive hyper | Repetitive | Repetitive hyper |
| --- | --- | --- | --- | --- |
| Number of A-to-I editing sites by RESIC | 23,194 | 12,496 | 1,225 | 895 |
| Shared with RADAR database full list (%)<sup>1</sup> | 75.4 | 65.4 | 27.0 | 30.7 |
| Novel sites, non-shared with RADAR database full list (%) | 24.6 | 34.6 | 73.0 | 69.3 |

<sup>1</sup>We considered sites as “shared with RADAR” if they were either included in the collection taken from RADAR, or their annotated genes are in RADAR’s collection.

Among all classes defined *via* RESIC, “non-repetitive” is the class yielded the largest overlap (75.4%, 65.4%, 27.0%, and 30.7%, for Nonrepetitive, Nonrepetitive hyper, Repetitive, and Repetitive hyper, respectively; **Table 1**). For repetitive site classes, smaller overlap was obtained.

As marked from our results, a substantial portion of sites detected by RESIC were not identified by others. The explanation for the new identified sites in this study may be the result of the usage of different tools for alignment, (i.e. Bowtie in our case and BWA, or a combination of Bowtie, SOAP, and BWA in (Zhu et al., 2013; Porath et al., 2014), as well as various threshold parameters and filtering criteria being set to consider sites as “editing sites” across tools.

The distribution of the editing events divided into classes can be shown in **Figure 2**, presenting for example the RESIC results for an adrenal tissue sample (the plots obtained from the rest of the samples can be found in the Supplementary Material (**Figure S3 -Figure S8**). A-to-G and T-to-C are both considered as editing changes because the data is not stranded. Over all classes being identified according to the graph alignment, A-to-G and T-to-C types were highly enriched, as expected (**Figure 2**). While non A-to-G mismatches are expected to be uncommon (Li et al., 2011; Kleinman and Majewski, 2012), RESIC still identified a certain amount of sites of that type, although in a much lower extent. This may be the result of rare SNPs that are uncovered by the SNP database being used.

**Figure 2:**
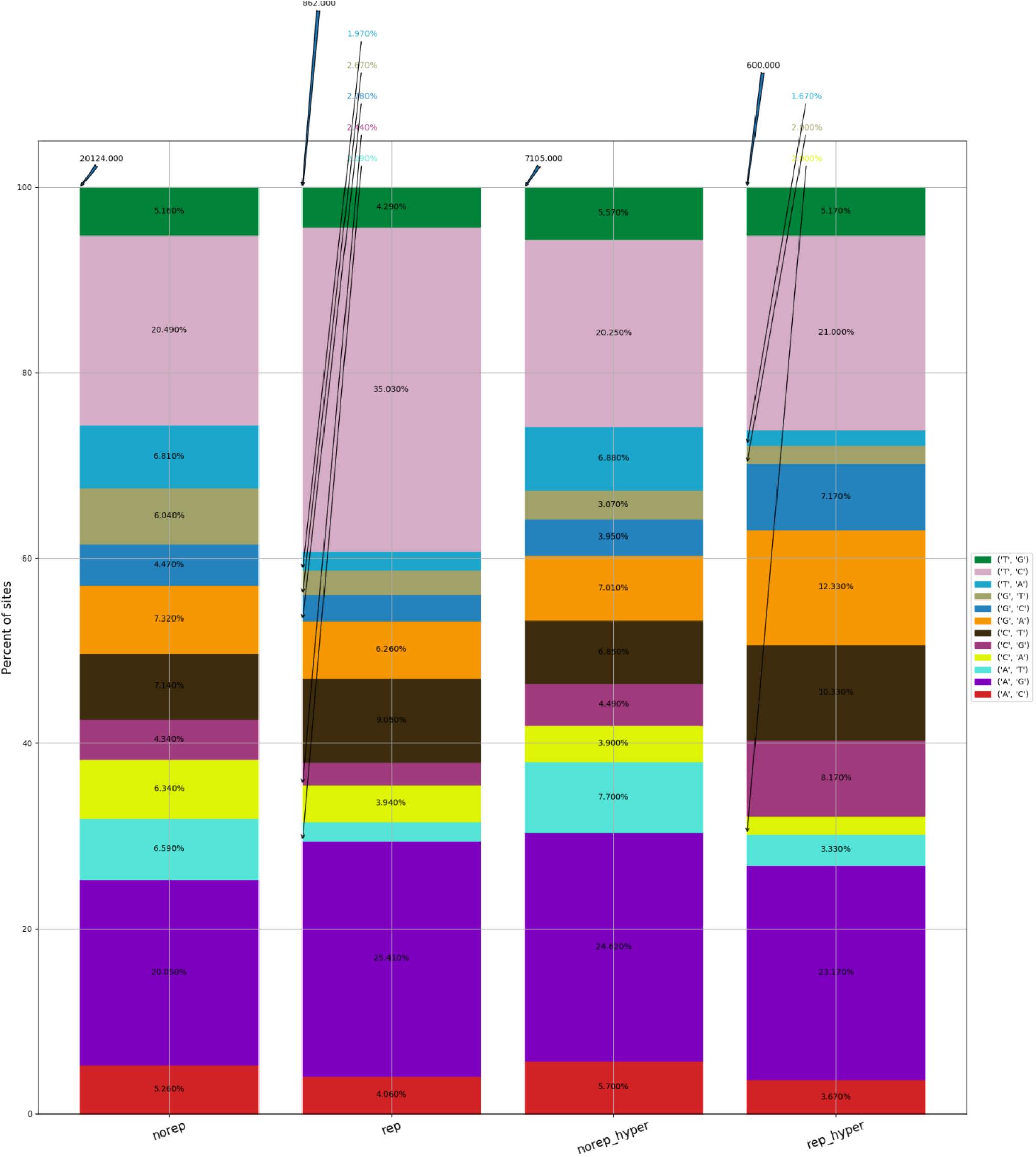
An example for RESIC editing percent distribution plot, obtained for an adrenal tissue sample. Blue arrow at the top of each bar shows the total number of sites being identified for the class. The percentages on the bars present the total number of editing type out of all identified site in the class.

Overall, the unique characterization of RESIC enables the detection of different classes of editing events, in one tool. The specificity of RESIC can be seamlessly controlled by modifying the running parameters, and by suppling datasets to exclude SNPs.

### SARS CoV-2 infection results in an extensive A-to-I hyper RNA editing and upregulation of ADAR1 enzyme

Systemic inflammatory responses to viral infection are triggered by IFN-mediated innate immune response (Schneider et al., 2014). Properly orchestrated, this type of immune response leads to inhibition of virus replication, promotion of virus clearance and induction of tissue repair. However, in some people infected with COVID-19, unpredictably, the innate immune response is exaggerated leading to Acute Respiratory Distress Syndrome (ARDS) (Moore and June, 2020). The innate immune response is regulated by ADAR enzymes, which modulate the interferon response to viral infection and reduce the innate immune response. ADAR1 was also shown to prevent MDA5 from sensing dsRNA (Liddicoat et al., 2015) and activating both type I and type III IFNs (Baños-Lara et al., 2013). Therefore, we wished to interrogate the activity of ADAR enzymes following SARS-CoV-2 infection. For this purpose, we analyzed the data of Blanco-Melo et al, of human Calu3 cells infected with SARS-CoV-2 virus or mock (Blanco-Melo et al., 2020), to examine the differences in A-to-I RNA editing patterns.

To identify and classify RNA editing sites, we applied RESIC on Calu3 cell lines that were infected with SARS-CoV-2 virus or mock (see Methods). Following RESIC run, we assessed the numbers of the most prevalent classes of A-to-I RNA editing types: non-repetitive, and hyper non-repetitive. We compared between SARS-CoV-2 and mock editing site numbers for each class following normalization, relying on the total read processed in each node. A-to-I non-repetitive hyper editing was much frequent in SARS-CoV-2 infected cells compared to mock (The number of hyper editing sites upon SARS-CoV-2 infection was 36.45% greater than in mock; **Figure 3**). In addition, although in the non-repetitive class the site numbers were relatively low for both sample types, the number of non-repetitive editing sites was larger as well in SARS-CoV-2 (The number of non-repetitive sites upon SARS-CoV-2 infection was 28.62% greater than in mock; **Figure 3**).

**Figure 3:**
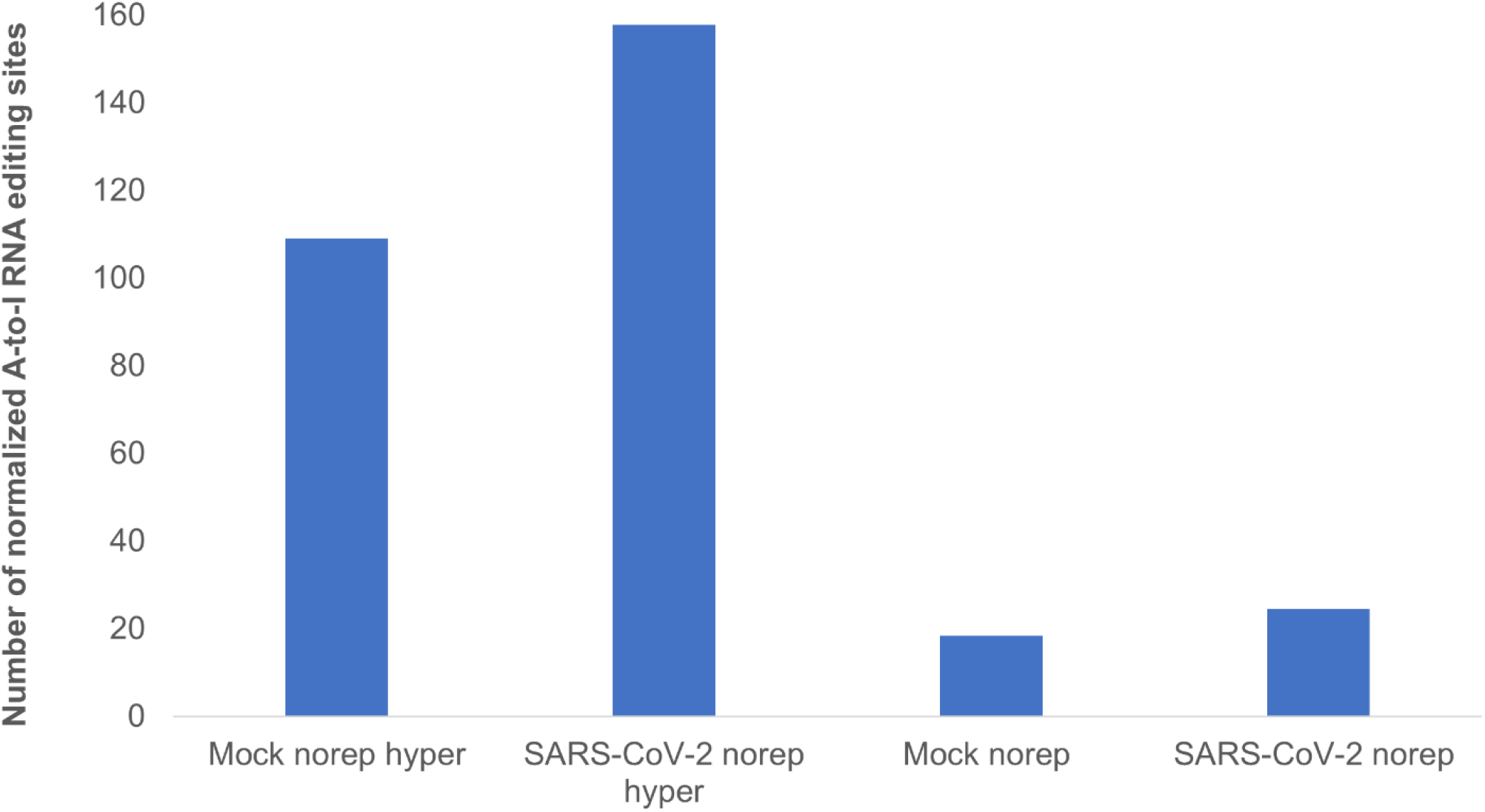
A-to-I hyper editing sites are more frequent in SARS-CoV-2 infected cells compared to mock. Presented here are A-to-I editing detected in SARS-CoV-2 or Mock samples in total, normalized to total counts. Nonrep: nonrepetitive sites, hyper: non-repetitive hyper editing sites.

To understand whether the higher editing activity upon SRAS-CoV-2 infection is manifested by a larger number of sites in the same genes as in mock, or additional sites located in new genes, we characterized the editing landscape with respect to site locations and gene annotation. We first tested the ratio of shared sites between SARS-CoV-2 and mock samples. For both non-repetitive and hyper non-repetitive classes, almost ∼70% of the sites were unique among samples infected with SARS-CoV-2, and about 60% of the sites were unique for mock samples (**Supplementary Table S2**). We next annotated the unique sites for each sample type, across different classes. It was apparent that the answer for our initial question is that among the unique editing sites following SARS-CoV-2 infection, some of the sites were extended the editing in baseline genes (considering the mock as the baseline), but most of the sites were in new genes (**Supplementary Table S2**).

Interestingly, the gene APOBEC3C became hyper edited following SARS-CoV-2 infection (**Supplementary Table S3**), while in mock samples it was classified under the non-repetitive class. The APOBEC family of enzymes edits C-to-U RNA modifications and known to be involved in regulation of innate immune response (Rosenberg et al., 2011; Schaefer et al., 2017). C-to-U editing of antibody-coding genes in the host’s DNA leads to diversification of the repertoire of antibodies produced against viruses, called somatic hypermutation (SHM) (Cogné, 2013). Therefore, hyper editing in APOBC genes may indicate for their involvement in COVID-19 phenotype, as part of a complex immune regulation system, controlling by A-to-I RNA editing. To test if the sets of unique edited genes display shared biological processes, we examined their biological process enrichment, using the web-based tool GeneMANIA (Zuberi et al., 2013). Submitting the list of hyper-nonrepetitive unique edited genes, upon SARS-CoV-2 infection, resulted in the enrichment of processes related to the regulation of I-kappaB kinase/NF-kappaB signaling (P-adjusted values of related pathways in the range of 1.91E-03 -3.50E-04; **Supplementary Table S3**). This result is intriguing in the light of strong indications suggesting that NF-kappaB pathway signaling has a critical role in controlling an excessive immune activation and ARDS (Kircheis et al., 2020). These indications, together with our result of hyper-editing in genes participating in the NF-kappaB pathway, suggest that A-to-I RNA editing activity may be critical to define the progression of COVID-19 disease and the risk to develop ARDS. We next, tested an enrichment for genes that were classified under the non-repetitive class and were uniquely edited in SARS-CoV-2 samples. Strong enrichment was obtained for processes related to interferon response (P-adjusted values of related pathways in the range of 1.25E-06 -1.03E-12; **Supplementary Table S4**), corroborating previous evidence that ADARs control interferon activation under viral infections (Baños-Lara et al., 2013), and suggests particularly that in COVID-19, ADARs control the level of immune response. We further run GeneMANIA for biological process enrichment with the mock unique gene sets, as a control. No processes related to interferon or NF-kappaB signaling were enriched in FDR<0.05 (**Tables S5-S6**).

Given these observations, we reasoned that interrogating the differences in ADAR RNA expression levels between SARS-CoV-2 and mock treated samples, would help to complete the picture. Therefore, we performed Differential Expression Analysis (DEA), on the same data of Blanco-Melo et al (Blanco-Melo et al., 2020) used for RESIC. We first created the read counts for both SARA-CoV-2 and mock Calu3 samples (see Methods). We than compared our created gene counts to those reported by Blanco-Melo et al (Blanco-Melo et al., 2020) GEO accession number GSE147507. Although read count values were not identical, as expected due to the use of different alignment tools, the trend was the same.

We performed DEA to identify changes in the expression of ADAR genes and A-to-I RNA editing, following SARS-CoV-2 infection (**Supplementary Table S7**). We found that all 10 ADAR1 isoforms are significantly upregulated **Figure 4**, P-value = 2.2e-16) in SARS-CoV-2 Calu3 infected cells. In comparison, the expression of ADAR2 did not differ between SARS-CoV-2 and mock samples (**Figure 4**, P-value >> 0.05). We also found that *IFIH1* (NM_022168.4), that encodes for Melanoma Differentiation-Associated Protein 5 (MDA5) is significantly up-regulated in the SARS-CoV-2 infected samples (**Supplementary Table S7**, P-adjusted = 7.46e-137). This is in line with findings indicating that the IFN response upon SARS-CoV-2 infection is primarily regulated by MDA5 (Yin et al., 2021). Since ADAR1 is known to prevent MDA5 from sensing dsRNA (Liddicoat et al., 2015), this result strengthening the conclusion that ADAR1 is largely involves in the immune response following SARS-CoV-2.

**Figure 4:**
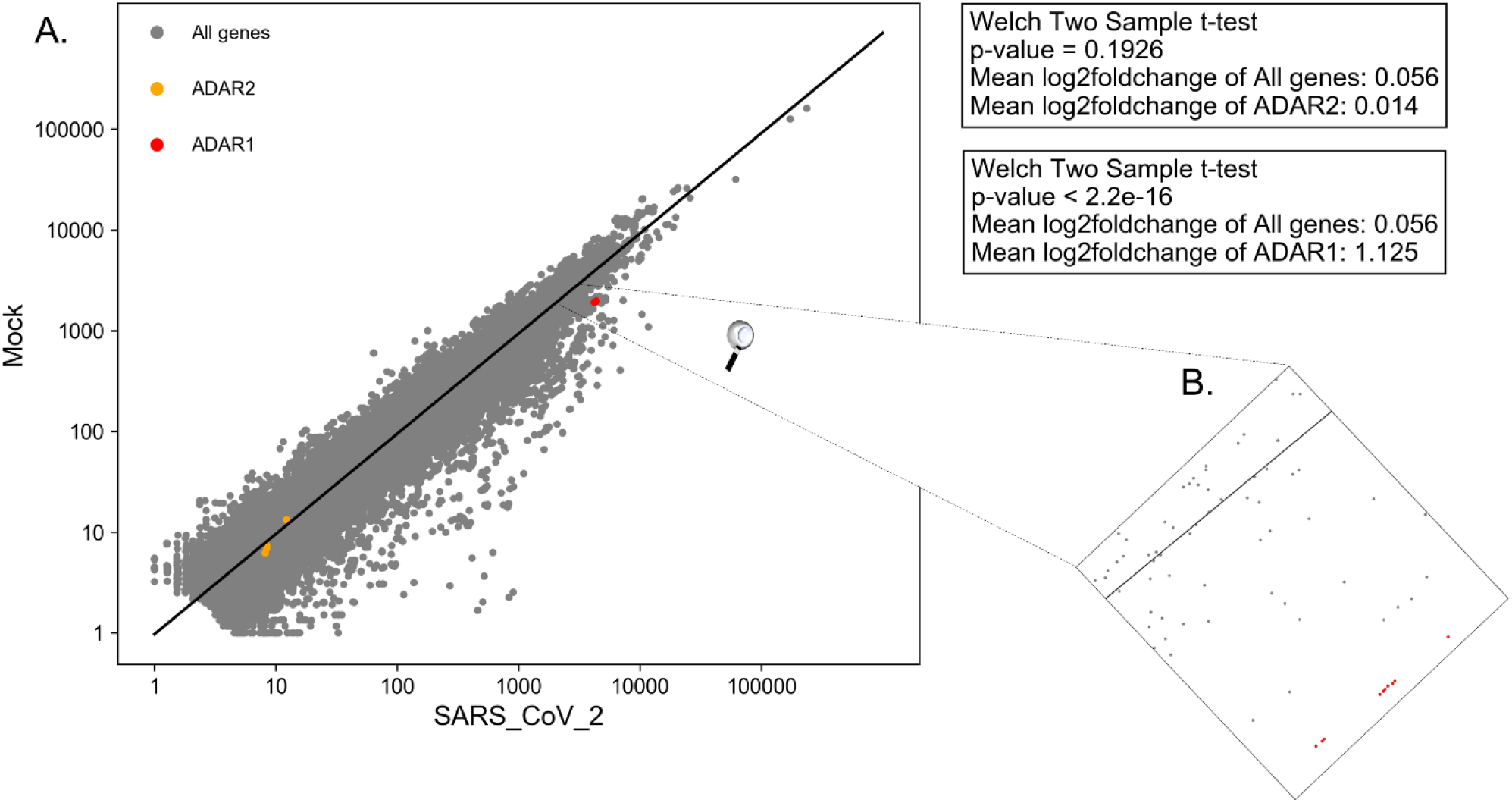
Significant upregulation of ADAR1 isoforms, but not ADAR2, in cells infected with SARS-CoV-2 virus, compared to mock. (**A**) Log scale plot shows normalized gene counts from mock cells against cells infected by SARS-CoV-2 virus. Every dot in the graph represents a gene: *ADAR1* isoforms (red), *ADAR2* isoforms (orange), and All other genes (grey). The black line is the regression line for all genes. The P-values were obtained using a Welch two-sample T-test on only transcripts with coefficient variation>1. **(B)** Zoom in on *ADAR1* isoforms on the log scale plot. All 10 isoforms are closely located on the plot.

Collectively, we suggest that upon SARS-CoV-2 infection, compared to mock 1) the number of hyper editing sites is increased; 2) ADAR1 activity is enhanced. The combination between these two observations goes together with the finding that ADAR1 is the enzyme mostly catalyzing hyper editing sites (Porath et al., 2014).

We tested if these results hold true for more *in-vitro* SARS-CoV-2 infected cell types created in the same study. For that purpose, we downloaded from GEO the already processed gene count data, for A549 and NHBE cells infected with SARS-CoV-2 high-multiplicity of infection (MOI). We chose downloading the already processed gene count data after validating for Calu3 cells that the gene count values created by us and the downloaded gene count from GEO (accession number GSE147507) are of the same trend (see Methods). For A549 cells, with a vector expressing human ACE2, indeed ADAR1, but not ADAR2, was significantly upregulated following SARS-CoV-2 infection, corroborating our previous results for Calu3 cells. However, for NHBE cells, both ADAR1 and ADAR2 were not significantly changed after SARS-CoV-2 infection (**Supplementary Table S8**). The non-significant upregulation of ADAR1 in the last case may be because of the different cell types used. In any event, we concluded that unlike ADAR2 the expression of ADAR1 is substantially different upon SARS-CoV-2, at least in some cell types.

Taken together, our results suggest that the catalyzation of hyper editing sites by ADAR1 is enhanced following SARS-CoV-2 infection. These results are intriguing in the context of ADAR1 role to block the interferon response, and particularly the role of hyper editing events to suppress the interferon induction (Vitali and Scadden, 2010). We hypothesize that editing levels might be indicative of the progression of COVID-19 disease and the risk to develop ARDS, as holds true in autoimmune diseases, due to the editing effect on the interferon response. Therefore, these results shed new light on the involvement of A-to-I RNA editing mechanism in COVID-19 disease. We note that supporting experimental validation is required to assess our conclusions. Our analysis encourages further exhaustive study of A-to-I RNA editing role in COVID-19 disease.

## Author contribution

D.L, R.H, and A.T.L. conceived and designed the study. D.L, R.H, MY, TH, and TB implemented and developed RESIC. R.H analyzed the data. A.T.L supervised the work. D.L, R.H, and A.T.L wrote the manuscript with input from all authors.

## Supplementary Figures

**Figure S1:**
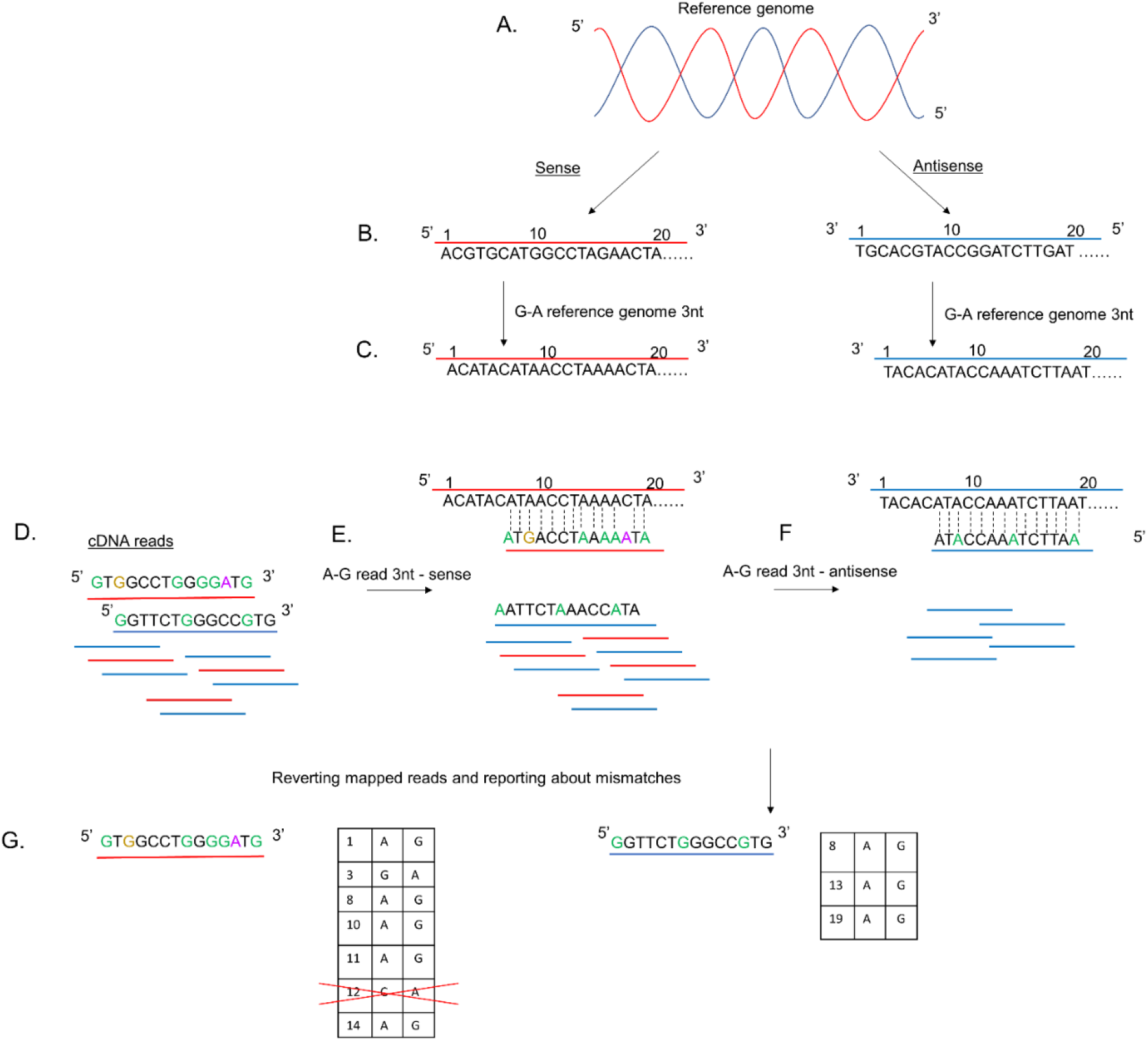
Detailed illustration of 3nt genome alignment scheme. **(A)** A reference genome. **(B)** The DNA sequence of the reference genome split by strand. Red, sense strand, presented in 5’-to-3’ direction; Blue, antisense strand, presented in 3’-to-5’ direction. The nucleotide sequences complement each other. **(C)** G-A 3nt replacement for the reference genome: Gs are converted into As **(D)** cDNA reads from RNA-sequencing, all obtained in 5’-to-3’ direction. Green nucleotides indicate for A-to-G editing sites (G instead of A in the reference genome). Brown nucleotides present G-to-A changes compared to the reference genome. Purple nucleotides present other mismatches than A-to-G or G-to-A. Red, sense strand; Blue, antisense strand. **(E)** Alignment following 3nt graph scheme -Part 1. read 3nt sense: Gs are converted into As in the cDNA reads. Hyper edited sense reads are successfully being mapped to the reference genome. **(F)** Alignment following 3nt graph scheme-Part 2, 3nt antisense: unmapped reads from part 1 are being reverted (not shown) and reversed to fit the 3’-to-5’ direction of the antisense reference strand. Gs are converted into As in the cDNA reads. Hyper edited antisense reads are successfully being mapped to the reference genome. **(G)** Reverting unmapped reads back from As to Gs and defining editing sites. Sites other than A-to-G or G-to-A are excluded.

**Figure S2:**
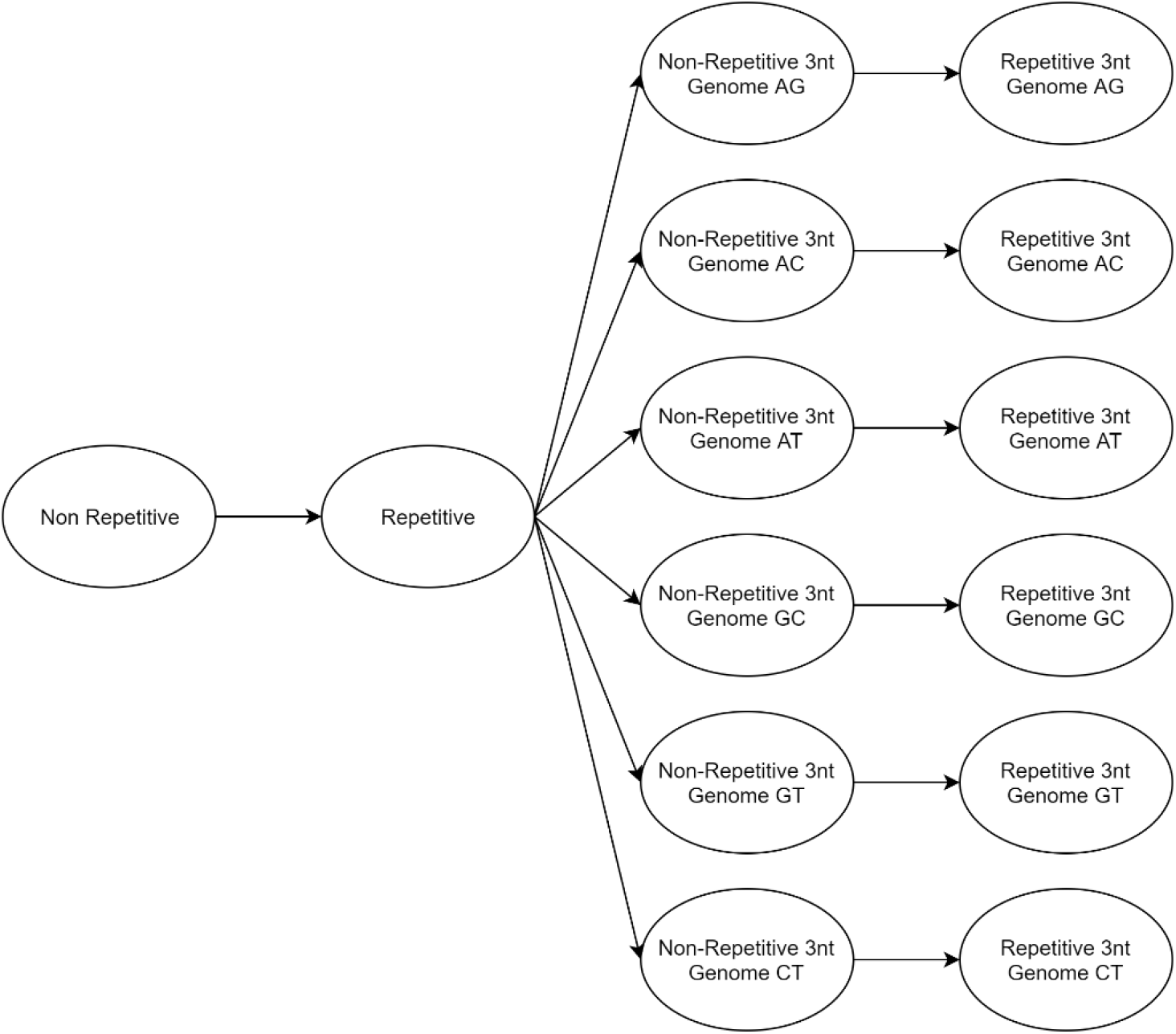
RESIC graph alignment. Non-repetitive alignment is followed by repetitive alignment. This is followed by 3nt Genome scheme with non-repetitive alignment parameters and 3nt Genome scheme with repetitive alignment parameters for each pair of nucleotides.

**Figure S3:**
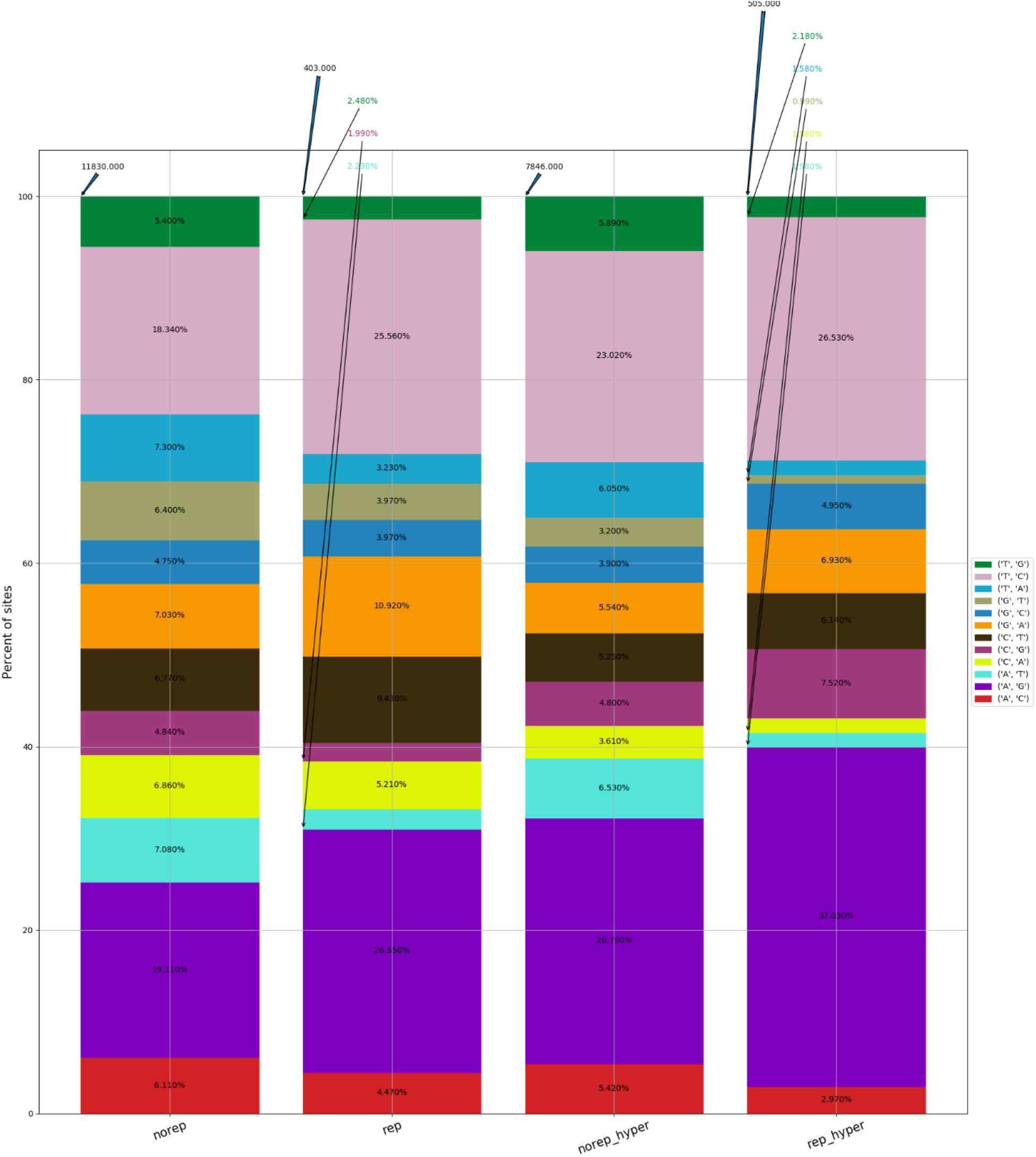
RESIC editing percent distribution plot, obtained for the adipose tissue. Blue arrow at the top of each bar shows the total number of sites being identified for the class. The percentages on the bars present the total number of editing type out of all identified site in the class.

**Figure S4:**
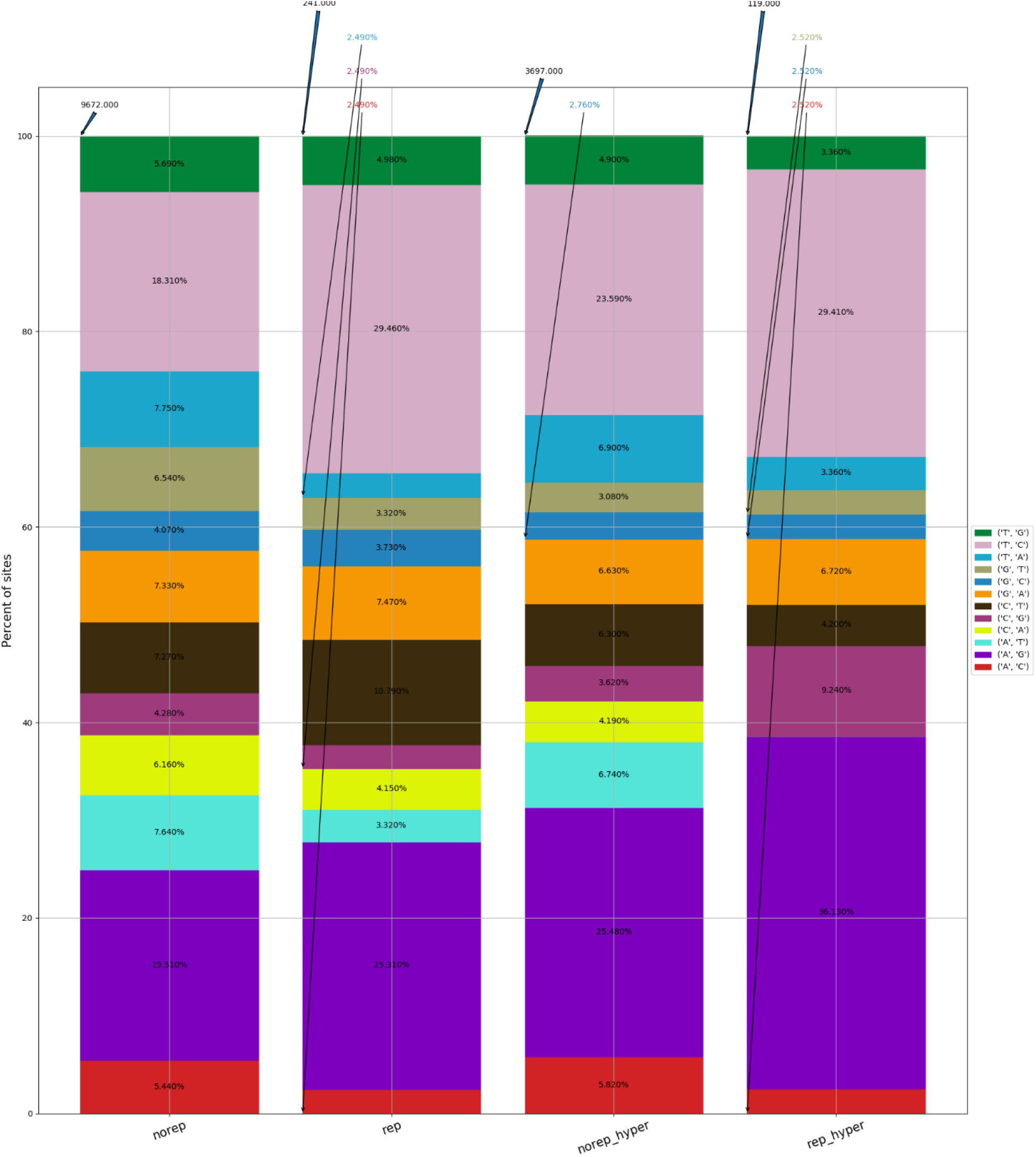
RESIC editing percent distribution plot, obtained for the brain tissue. Blue arrow at the top of each bar shows the total number of sites being identified for the class. The percentages on the bars present the total number of editing type out of all identified site in the class.

**FigureS5:**
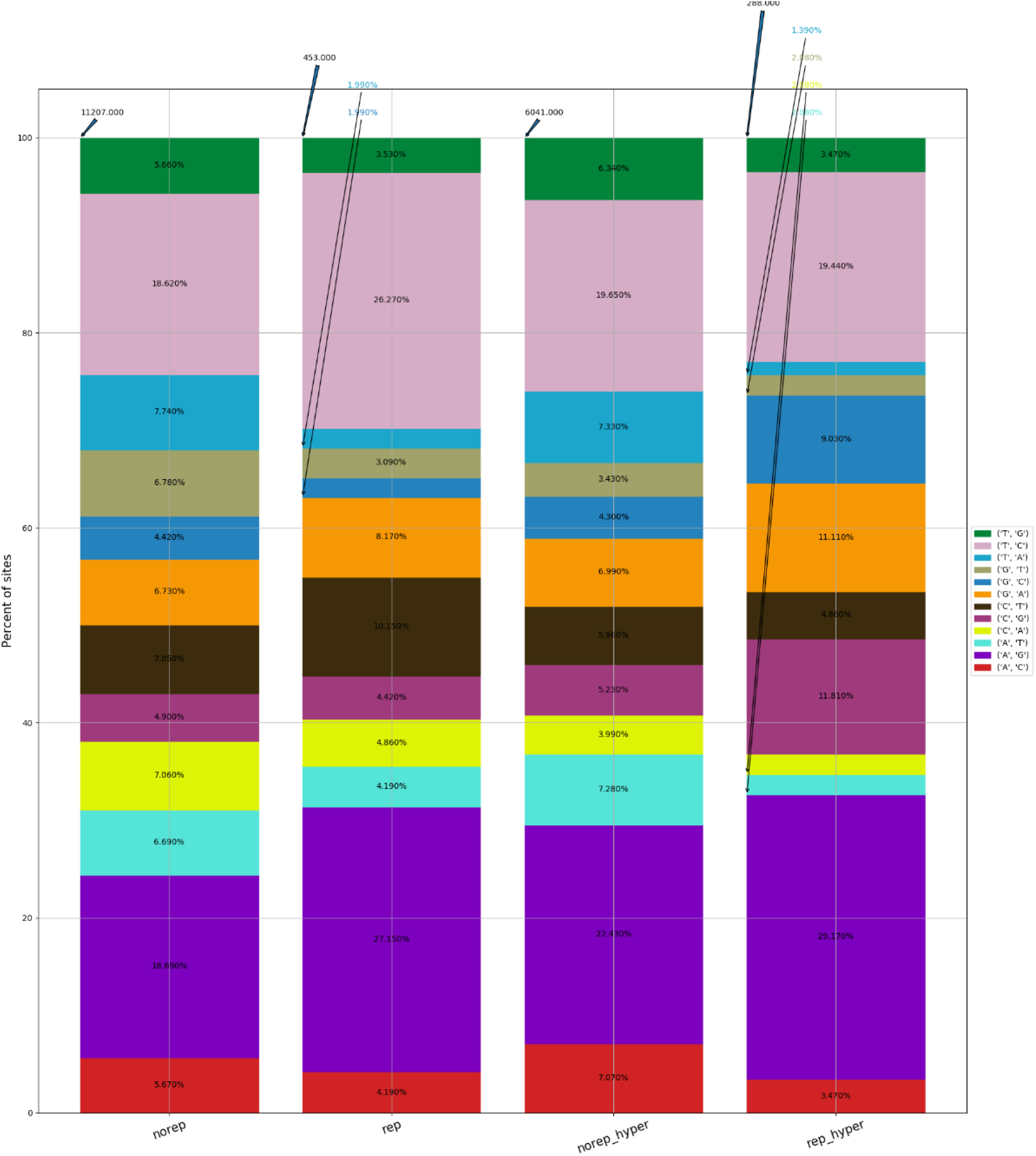
RESIC editing percent distribution plot, obtained for the breast tissue. Blue arrow at the top of each bar shows the total number of sites being identified for the class. The percentages on the bars present the total number of editing type out of all identified site in the class.

**FigureS6:**
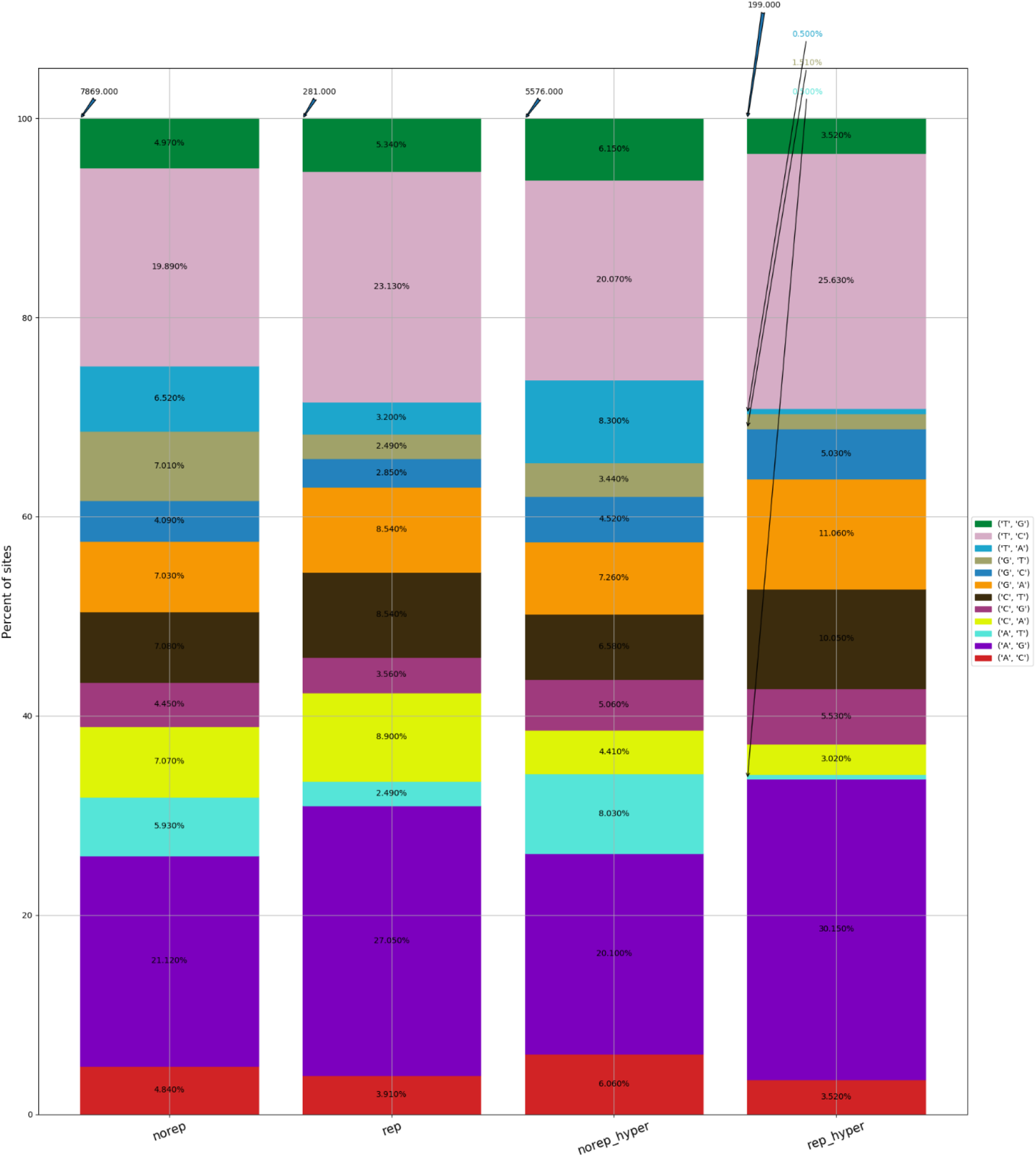
RESIC editing percent distribution plot, obtained for the colon tissue. Blue arrow at the top of each bar shows the total number of sites being identified for the class. The percentages on the bars present the total number of editing type out of all identified site in the class.

**FigureS7:**
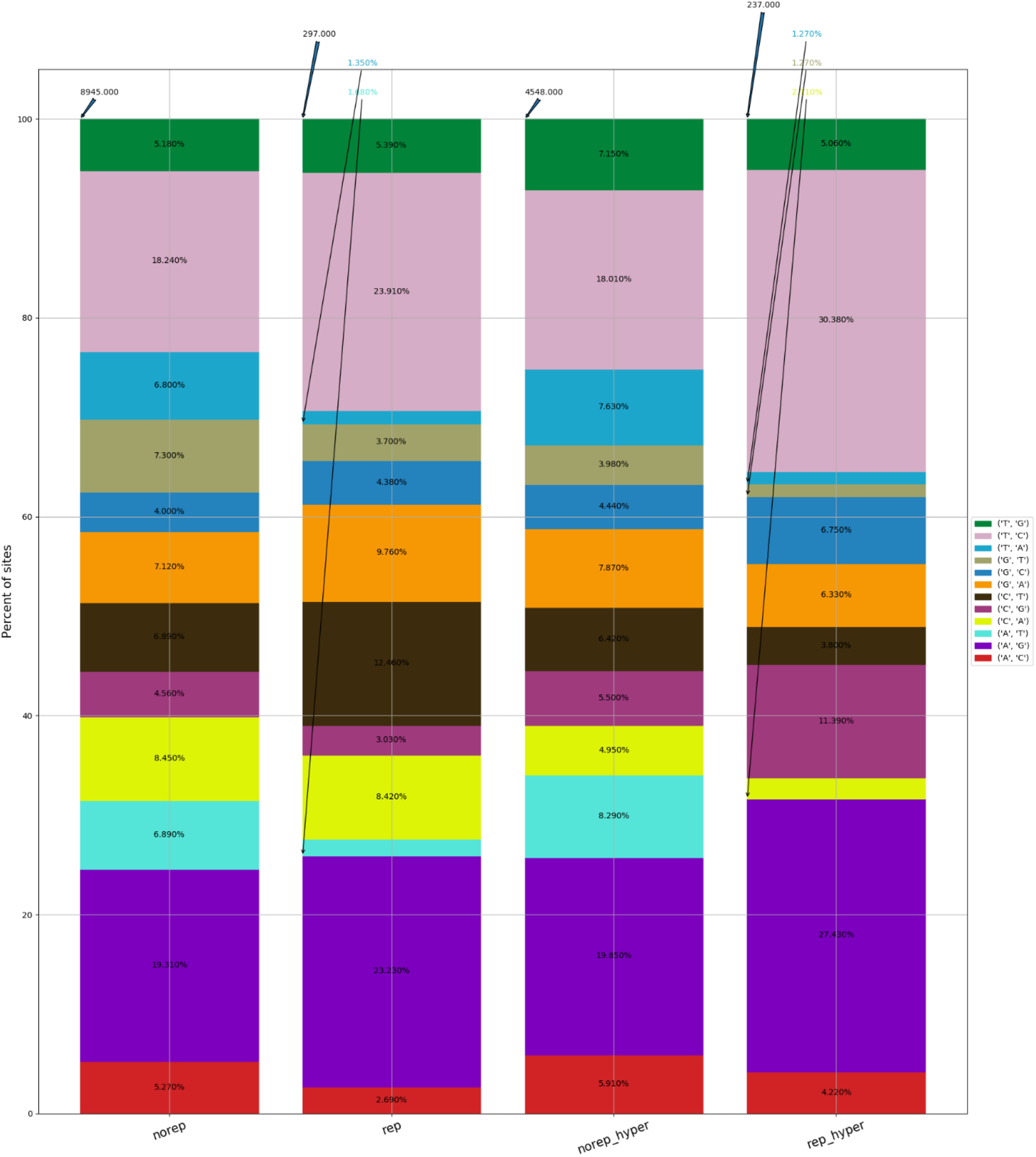
RESIC editing percent distribution plot, obtained for the kidney tissue. Blue arrow at the top of each bar shows the total number of sites being identified for the class. The percentages on the bars present the total number of editing type out of all identified site in the class.

**FigureS8:**
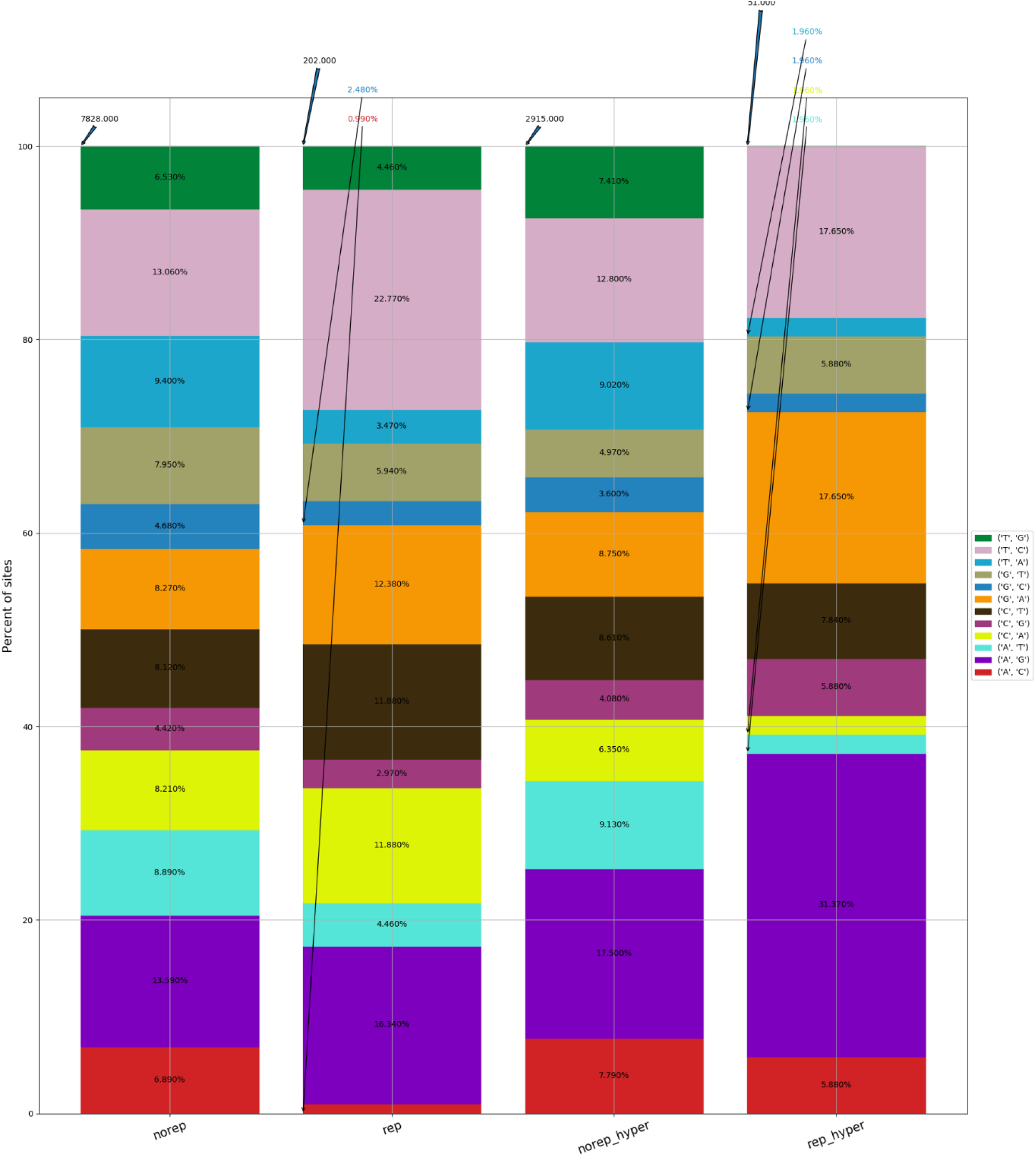
RESIC editing percent distribution plot, obtained for the heart tissue. Blue arrow at the top of each bar shows the total number of sites being identified for the class. The percentages on the bars present the total number of editing type out of all identified site in the class.

## Supplementary table legend

Supplementary Table S1: the list of editing sites across different classes identified using samples from the Illumina Human Body Map project.

Supplementary Table S2: characterization of editing landscape with respect to site locations and gene annotation

Supplementary Table S3: unique genes under the SARS-CoV-2 nonrepetitive hyper-editing class and GO enrichment analysis.

Supplementary Table S4: unique genes under the SARS-CoV-2 nonrepetitive class and GO enrichment analysis.

Supplementary Table S5: unique genes under the mock nonrepetitive hyper-editing class and GO enrichment analysis.

Supplementary Table S6: unique genes under the mock nonrepetitive class and GO enrichment analysis.

Supplementary Table S7: differential expression analysis for SARS-CoV-2 infected samples versus mock, in the Calu3 cell line.

Supplementary Table S8: differential expression results of ADAR1 and ADAR2 genes, for SARS-CoV-2 infected samples versus mock, in A549 and NHBE cell lines.

